# *Angelica keiskei* leaves extract attenuates psychosocial stress in overcrowding-subjected rats

**DOI:** 10.1101/2024.04.24.591022

**Authors:** Ferbian Milas Siswanto

## Abstract

*Angelica keiskei* is currently used as a popular functional food with various beneficial effects, including antioxidant, anti-obesity, anti-tumor, anti-diabetic, and anti-bacterial. A recent *in vitro* study reported that *A. keiskei* inhibits monoamine oxidases (MAOs), suggesting the antidepressant property of *A. keiskei*. However, *in vivo* studies on laboratory animals subjected to psychosocial stress have not been conducted. In this study, the effects of an *A. keiskei* leaves extract (AKE) on rats undergoing chronic social overcrowding stress were explored. Six-month-old male and female Wistar rats were housed in groups of three in 12 × 12 × 18 cm cages (overcrowding) for twenty-eight days. Both male and female rats were divided into two groups (N = 10); control group received oral distilled water, whilst the other group (treatment group) received 20 mg/kgBW/day of AKE supplementation. The results showed that AKE-treated rats exhibited lower anxiety- and depressive-related behaviors than that of control stressed rats. AKE significantly decreased corticosterone and increased testosterone and estrogen levels in stressed rats. Additionally, brain tissue MDA and TNF-alpha levels were reduced, while brain neurotransmitter 5-HT and antioxidant SOD levels were elevated by the AKE. These findings suggest, for the first time, that AKE could alleviate overcrowding stress-induced behavioral, neuroendocrine, antioxidant, and inflammatory dysfunctions. *A. keiskei* leaves extract could potentially be used as an agonist to reduce stress and depression.

**GRAPHICAL ABSTRACT:** 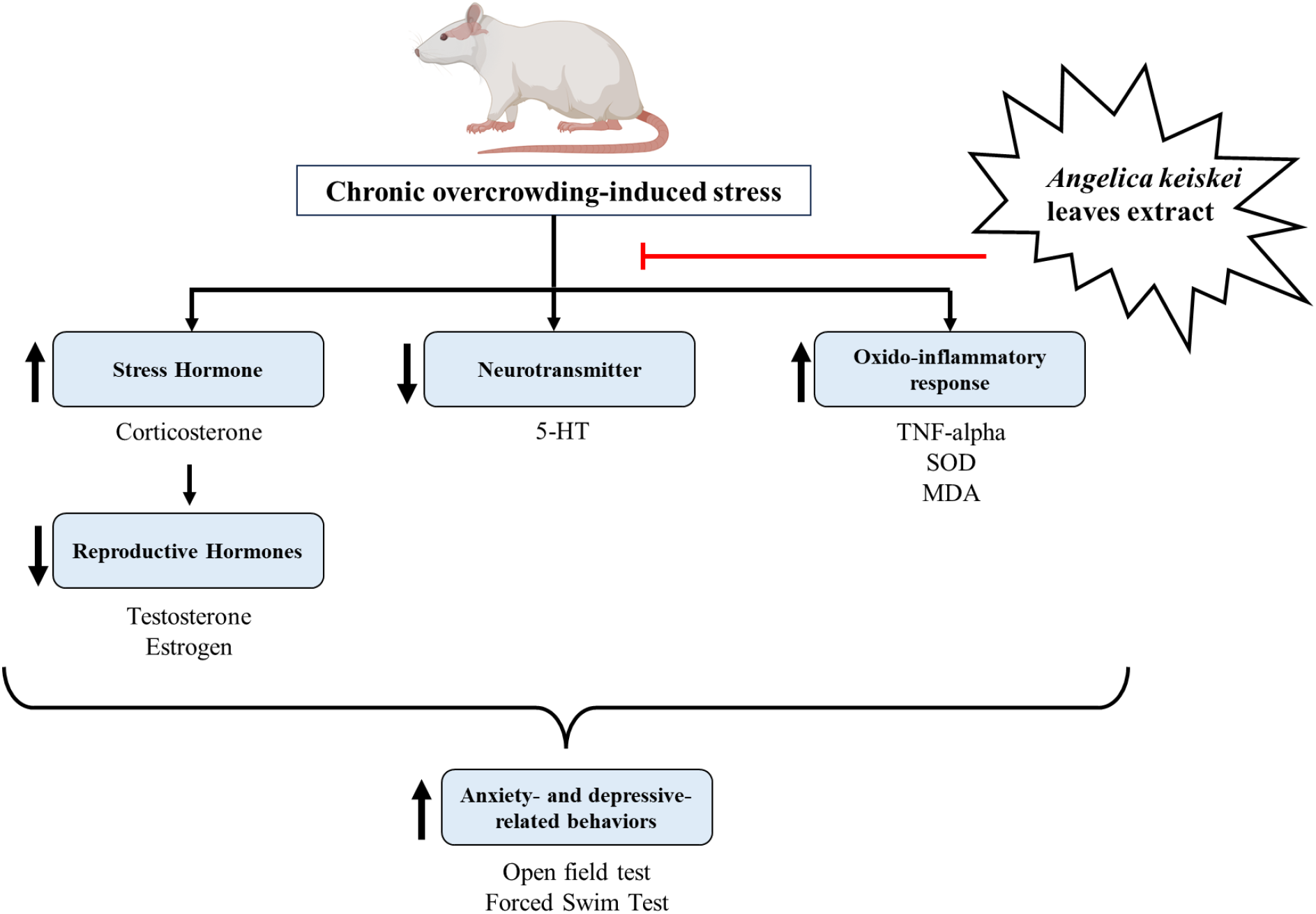

## I. INTRODUCTION

Stress is present in everyday life and is one of the most prevalent conditions experienced by humans, regardless of age, gender, race, and social status. Stress is defined as an internal perception towards real or perceived external stimuli that threatens homeostasis and triggers physiological responses which involves the endocrine, nervous, and immune systems [1,2]. Two stressors frequently investigated are physiological and psychosocial stresses. In addition to subjective and hormonal responses, both stresses also share common neural substrates [3]. Studies on animals as well as population-based research on humans have demonstrated the impact of chronic stress on metabolic, endocrine, and psychological processes. Chronic stress is linked to a higher risk of biological, psychological, and social issues. Moreover, stress is thought to be a significant risk factor for the aging process, and it can manifest early in life if it is persistently present [4].

Prolonged exposure to psychosocial stress has been linked to an increased incidence of major depressive disorder [5]. According to several animal model studies, several stressors can change the structure and function of the brain, resulting in behaviors linked to depression [6]. Psychosocial stress elicits biological and behavioral responses due to the activation of the hypothalamic-pituitary-adrenal (HPA) axis, particularly stress hormones known as corticosterone [7]. The HPA axis mediates the inhibition of the hypothalamic–pituitary– gonadal (HPG) axis, which is responsible for reproductive function and healthy aging in both men and women [8,9]. Additionally, studies showed that and increasing level of brain pro-inflammatory cytokines contributes to the pathophysiology of stress-induced anxiety and depression, suggesting the critical involvement of inflammation [10,11]. Oxidative stress, an imbalance between the production of radicals and antioxidants, also contributes to pathological conditions related to psychosocial stress [12]. There are several methods to induce psychosocial stress in experimental animals, including overcrowding, social defeat, the restrain stress model, etc. In this study, psychosocial stress is induced by the overcrowding method, which has been shown to effectively induce psychosocial stress in mice and rats and increase corticosterone [13,14].

To attenuate the negative impact of psychosocial stress, many investigations on the effectiveness and underlying processes of plant-based antidepressant and anti-stress medicines have been carried out recently. It is well documented that herbal medicines might lessen the negative impacts of anxiety and depression [15]. An *in vitro* study revealed that *Angelica keiskei*, a flavonoid-rich traditional plant from Japan, contains chemicals that selectively inhibit monoamine oxidases (MAOs) with an IC50 value of 3.43 μM, which suggests its antidepressant property [16]. Studies in mice with psychosocial stress have reported that flavonoid can prevent HPA axis activation, reduce corticosterone levels, regulate hippocampal glucocorticoid receptors (GR), and prevent nerve cell damage due to its activity to bind 5-HT1A [17]. In addition, a study reported that flavonoid inhibits corticosterone elevation due to reduced corticotropin-releasing factor (CRF) by blocking the DNA binding activities of the glucocorticoid receptor [18]. The preliminary study that has been conducted showed that 20 and 40 mg/kgBW/day of ethanolic extract of *Angelica keiskei* leaves (AKE) could prevent increased corticosterone levels in overcrowding-subjected male and female rats (data not shown).

Despite the broad effect of stress in human and *in vitro* studies that suggested the potential role of AKE to ameliorate stress-induced impairment, there has been no research focusing on the effect of AKE on overcrowding-induced stress in animal models, particularly the impact on behavioral, neuroendocrine, antioxidant, and inflammatory responses. Thus, this study aimed to analyze the effects of AKE in overcrowding-subjected rats and focused on the application of AKE to improve several behavioral (anxiety- and depressive-like behaviors), neuroendocrine (5-HT, corticosterone, testosterone, and estrogen), antioxidant (SOD and MDA), and inflammatory (TNF-alpha) parameters, explicitly emphasizing the differential effects of AKE on male and female rats.

## 2. MATERIALS AND METHODS

### 2.1. Animals and experimental groups

This study used a randomized pretest-posttest control group design, carried out using health rats (*Rattus norvegicus*), Wistar strain, male (n *=* 20) and female (n *=* 20), healthy, aged 6 months, weighing 180–200 grams. The rats were purchased from the experimental animal breeding facility of the Medical Faculty, Udayana University, and were kept under standard conditions [19]. The environmental conditions for animal housing were controlled at stable condition of 12 hr light/dark cycle, reduced light intensity (1 - 25 lx), temperature of 22 – 24 °C, humidity of 40-60%, and ad libitum sources of food and water throughout the experiment. Male and female rats were each divided randomly into two groups (10 rats per group). The control group was given 2 ml of distilled water orally two hours prior to psychosocial stress induction. The treatment group was given 20 mg/kgBW/day of AKE wo hours prior to psychosocial stress induction. Psychosocial stress was induced using the previously reported overcrowding method [13]. In brief, rats were placed in 12 cm × 12 cm × 18 cm cages, three rats per cage (movement space per rat was 48 cm^2^), for four hours per day, twenty-eight days in a row. After four hours, rats were returned to their respective standard 40 cm × 30 cm × 15 cm cages (3 rats per cage, movement space per rat was 400 cm^2^). All experiments were conducted according to the instructions of the Ethical Committee of the Faculty of Medicine and Health Sciences, Atma Jaya Catholic University of Indonesia.

### 2.2. Preparation of the extract

Dried leaves of *Angelica keiskei* were purchased from a local market (Denpasar, Bali, Indonesia), ground into a coarse powder, and passed through a 40-mesh sieve (425 μm). *Angelica keiskei* leaves powder was mixed with five volumes of 70% ethanol for 48 hours, filtered using filter paper (Whatman No. 1), and the solvent was dissolved with a rotary evaporator. Then, the ethanolic extract of *Angelica keiskei* leaves (AKE) was kept at −20°C until further use.

### 2.3. Behavioral tests

Before and after twenty-eight days of overcrowding and AKE treatment, rats were subjected to behavioral testing, including an open field test and a forced swim test to assess the anxiety- and depressive-related behaviors, respectively. All behavior tests were carried out in the light phase. In the open field test, rats were placed individually in the center of an acrylic box (40 cm × 40 cm × 40 cm). The base of the box was equipped with 10 × 10 cm square line grids. The movements of each rat were recorded for six minutes. The time spent in the four squares in the center of grids and twelve peripheral squares was calculated. Next, the rearing behavior, or the number of times the rat completely stood on both legs in a vertical upright position, was measured in the open field test as an indicator of anxiety level. The chamber was wiped with 95% ethanol between trials to remove any scent clues left by the previous subject rat [20]. In the forced swim test, rats were individually placed in transparent cylindrical glass containers (21 cm diameter, 50 cm height) containing 23 ºC water for 5 min. After 5 minutes, rats were removed to a transient drying cage. The forced swim test was recorded, and the amount of time rats remained immobile was quantified as depressive-like behavior [21].

### 2.4. Hormonal analysis

The levels of corticosterone, testosterone, and estrogen were quantified from rats’ serum. Blood was collected before and twenty-eight days after treatment by utilizing sterile 75 μl EDTA-coated capillary tubes to puncture the retro-bulbar sinus from the medial canthus of the eye, and serum was isolated by centrifugation. The levels of serum corticosterone, testosterone, and estrogen were measured by enzyme-linked immunosorbent assay (ELISA) kits purchased from Bioassay Technology Laboratory (Shanghai, China) following the manufacturer’s instructions for corticosterone (Cat no. E0828Ra), testosterone (Cat no. EA0023Ra), and estrogen (Cat no. E1297Ra).

### 2.5. Measurement of brain tissue 5-HT, SOD, MDA, and TNF-alpha levels

The levels of neurotransmitter 5-hydroxytryptamine (5-HT), oxidative stress markers (SOD and MDA), and inflammatory cytokines (TNF-alpha) were measured from the brain tissue (prefrontal cortex) homogenates after behavioral tests and blood collection. To obtain brain tissue homogenates, the brains of the rats were collected and homogenized in Dulbecco’s Modified Eagle Medium (DMEM) F-12 (Wako Pure Chemical, Osaka, Japan) on ice, centrifuged at 4°C for 20 minutes at 10,000 rpm, and the obtained supernatant was collected for further experiments. The levels of 5-HT (Cat no. CSB-E08364r, Cusabio, Texas, USA), SOD (Cat no. E1444Ra, Bioassay Technology Laboratory), MDA (Cat no. E0156Ra, Bioassay Technology Laboratory), and TNF-alpha (Cat no. E0764Ra, Bioassay Technology Laboratory) were measured in the brain tissue homogenates using commercially available rat ELISA kits according to the respective manufacturer’s protocol.

### 2.6. Data Analysis

The data were presented as mean ± standard error of the mean (SEM) and analyzed with IBM SPSS Statistics for Windows, version 23.0. Statistical comparisons were performed with independent sample *t*-test or mixed analysis of variance (ANOVA) as described in the figure legends. Differences were significant when p <0.05 (*), <0.01 (**), or <0.001 (***).

## 3. RESULTS

### 3.1. AKE attenuates anxiety- and depressive-like behaviors caused by overcrowding

In the open field test, chronic overcrowding in both male and female control rats significantly reduced the time spent in the center grids (p < 0.001). Inversely, the total duration of time spent in the peripheral grids was higher in the overcrowding-induced stress groups, both male and female rats (p < 0.001). Additionally, overcrowding significantly reduced the number of rearing movements in both males and females to a similar extent (p < 0.001). Together, these results suggest that overcrowding induces anxiety-like behavior to a similar degree in both males and females. Stress led to a considerable increment in the duration of immobilization in the forced swim test (p < 0.001), and this elevation was higher in female rats (131.00 ± 10.66 s) compared to male rats (111.57 ± 10.16 s). These results indicate that female rats show more depressive-like behavior than male rats following overcrowding-induced stress. Next, the effects of AKE on these anxiety- and depressive-like behaviors were tested. Daily oral administration of AKE for twenty-eight days attenuated the reduced time spent in the center grids (p < 0.001 for both male and female), increased time spent in the peripheral grids (p < 0.01 for male and p < 0.001 for female), reduced number of rearing movements (p < 0.001 for male and p < 0.05 for female), and increased time of immobilization (p < 0.001 for both male and female). Together, these findings suggest that AKE presents antidepressant activity in animal models of anxiety and depression (Figure 1). In addition, any adverse effect on the group of rats treated with AKE was not observed at any time point.

**Figure 1.**
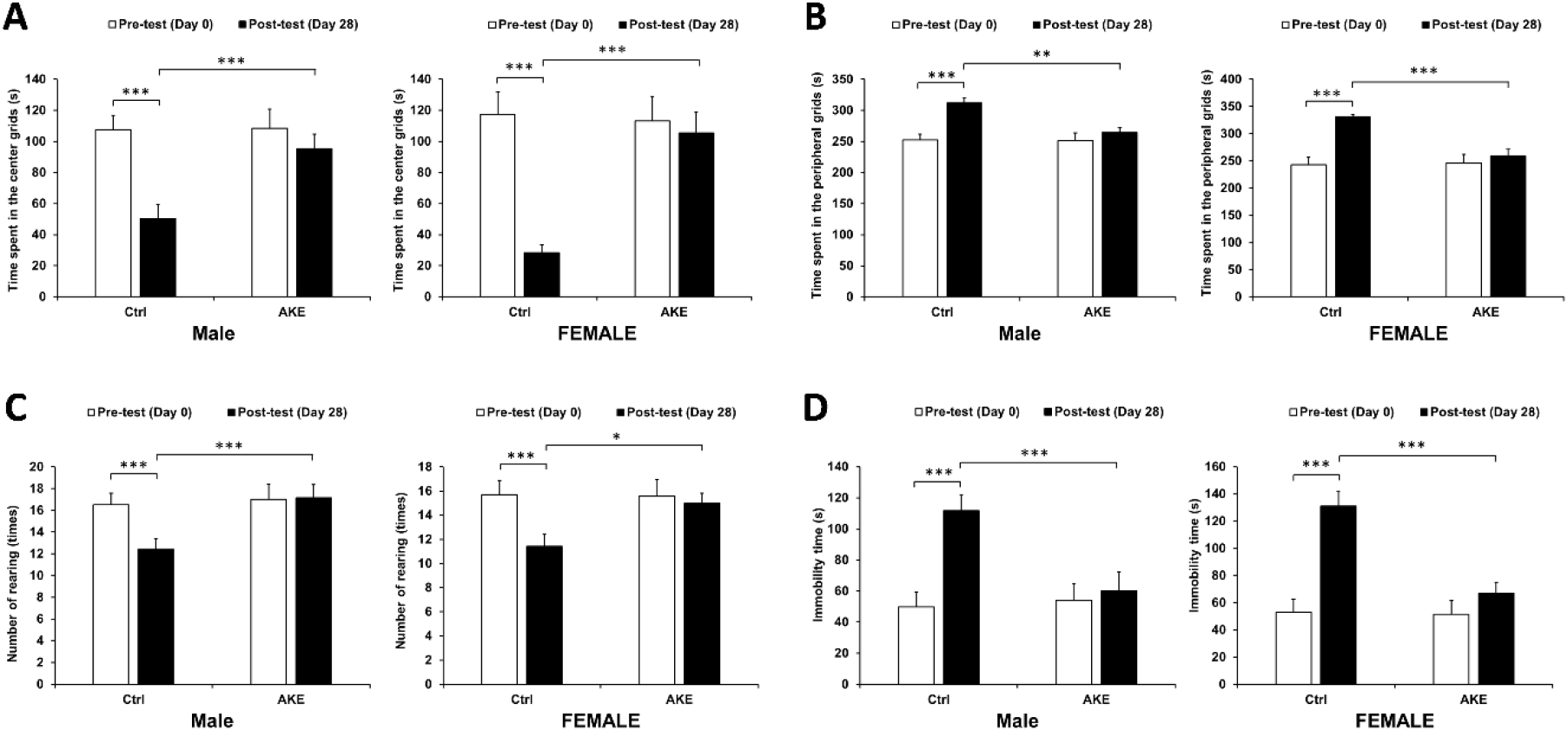
Effects of overcrowding and AKE on anxiety- and depressive-related behaviors. (A-C) AKE-treated stressed male and female rats exhibited a lower anxiety-like behavior than that of control stressed rats in the open field test, as shown by reduced time spent in the center grids (A), increased time spent in the peripheral grids (B), and a reduced number of forelimbs rearing (C). (D) AKE-treated stressed male and female rats exhibited a lower depressive-like behavior than that of control stressed rats in the forced swim test, as indicated by increased time immobility. Data represent means ± SEM (n = 10), mixed ANOVA, *p<0.05, **p<0.01, ***p<0.001.

#### 3.2. Effects of AKE on the hormonal biomarkers in social overcrowding-subjected rats

Serum corticosterone (male and female), testosteron (male only), and estrogen (female only) levels were determined before and at the end of the experiment using the ELISA technique. Consistent with the anxiety- and depressive-like behavior, the exposure of rats to a twenty-eight-day of social overcrowding stress significantly raised the level of corticosterone in the control male (from 4.32 ± 0.90 to 18.01 ± 2.54 ng/mL, p < 0.001) and female (from 55.23 ± 10.23 to 191.42 ± 22.66 ng/mL, p < 0.001) rats. However, this increment was completely ameliorated by treatment with AKE (all p < 0.001). Next, the levels of testosterone and estrogen in male and female rats, respectively, as a representative of HPG axis function were measured. As expected, the testosterone level was significantly decreased by overcrowding (p < 0.001), and this depletion was partially restored by AKE treatment (p < 0.001). Similarly, estrogen levels were substantially declined by overcrowding (p < 0.001), and AKE protected rats from this depletion (p < 0.001). These findings suggest the beneficial effects of AKE on overcrowding-induced stress that may be attributed to blunting the detrimental effect of the stress on fertility (Figure 2).

**Figure 2.**
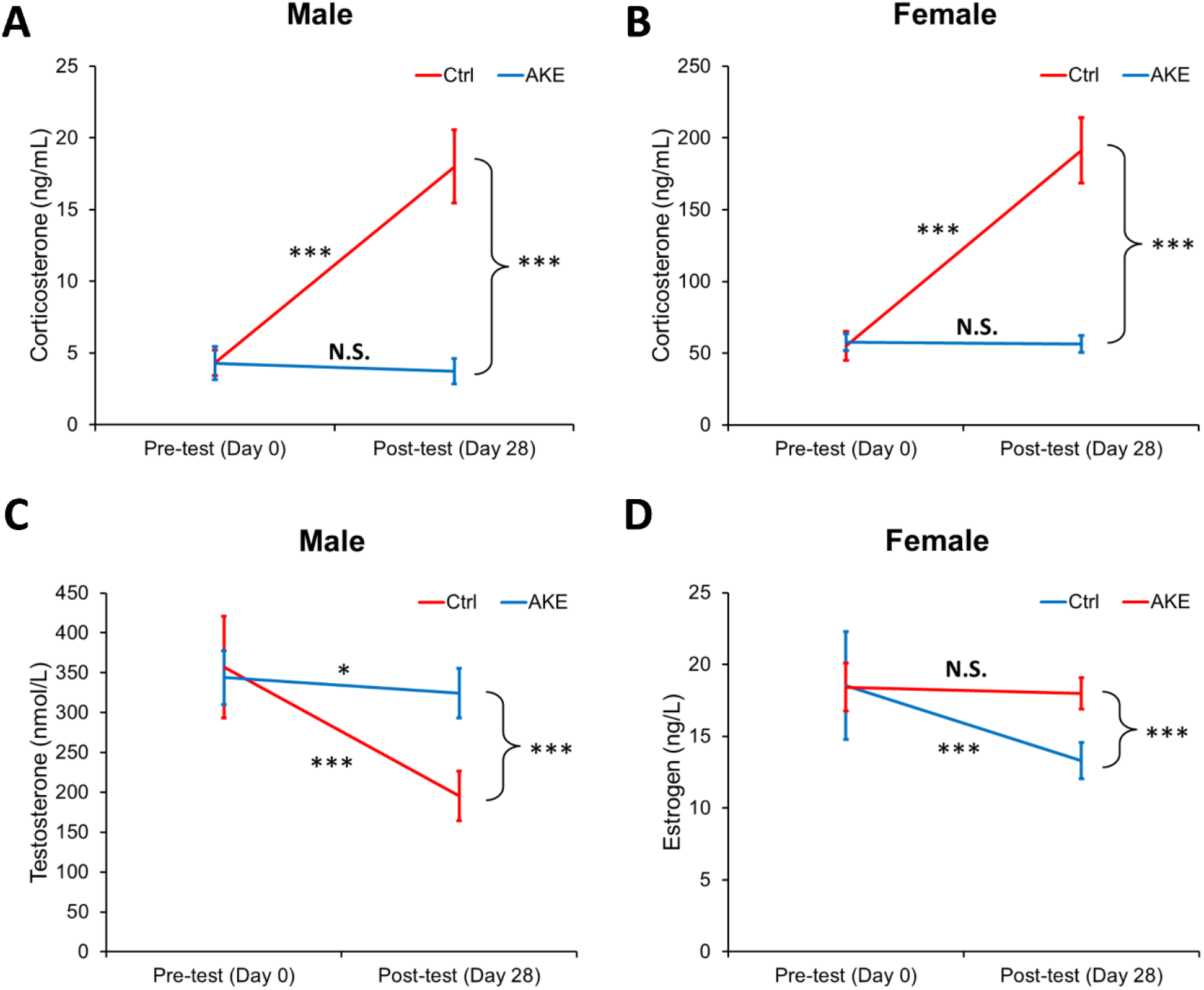
Changes in plasma corticosterone, testosterone, and estrogen levels in overcrowding- and AKE-treated rats. (A) The plasma concentration of corticosterone in male rats; (B) the plasma concentration of corticosterone in female rats; (C) the plasma concentration of testosterone in male rats; and (D) the plasma concentration of estrogen in female rats before and after overcrowding and AKE treatment for twenty-eight days. Data represent means ± SEM (n = 10), mixed ANOVA, *p<0.05, **p<0.01, ***p<0.001.

### 3.3. Role of AKE on the expression of neurotransmitters in the brain

Psychogenic stressors have been widely established to affect the activity of neurotransmitters, the signaling molecules between neurons, particularly norepinephrine (NE), dopamine (DA), and serotonin (5-HT). Neurotransmitters are important for depression and stress [22]. In the present study, the levels of brain 5-HT as a representative monoamine neurotransmitter were determined at the end of the experiment using the ELISA technique. The levels of 5-HT were comparable in male and female rats. The AKE-treated group significantly elevated the brain 5-HT levels in stressed rats, both male and female (p < 0.001 for all) compared to the control stressed rats (Figure 3A). These results indicate that AKE restored the social overcrowding-induced depletion of monoamine neurotransmitter levels, which may play a beneficial role in the improvement of depression in rats.

**Figure 3.**
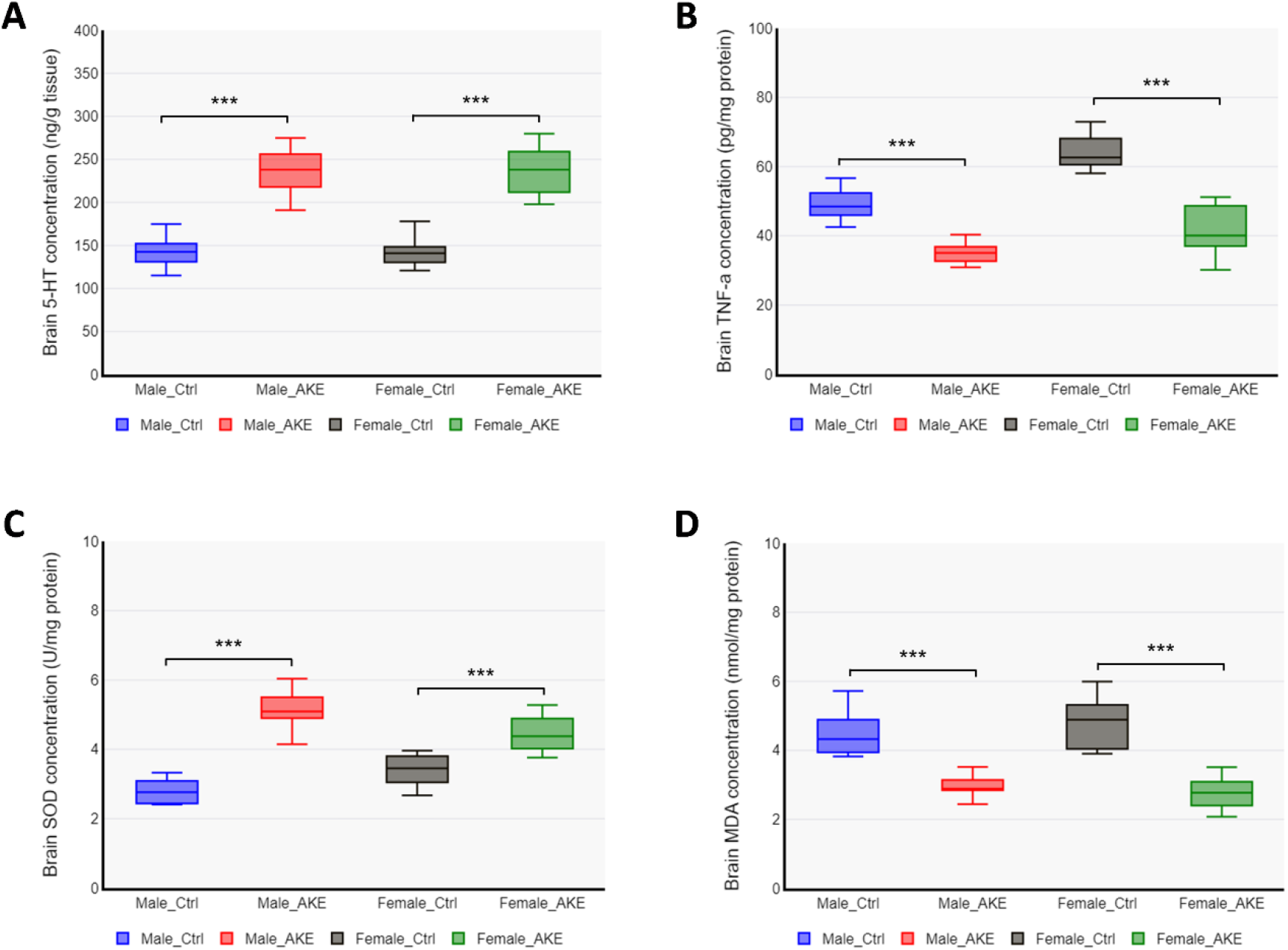
The effect of AKE on serotonin, TNF-alpha, and redox status biomarkers. Rats were subjected to overcrowding treatment for four hours for twenty-eight consecutive days, with or without AKE supplementation. (A) The neurotransmitter 5-HT, (B) the pro-inflammatory cytokine TNF-a, (C) the antioxidant enzyme SOD, and (D) the biomarker for oxidative damage MDA, were measured by ELISA from brain tissue homogenates. Data represent means ± SEM (n = 10), independent sample *t*-test, *p<0.05, **p<0.01, ***p<0.001.

### 3.4. AKE modulates the oxido-inflammatory responses of the brain

Depressive disorder caused by psychosocial stress is a combination of brain tissue damage as a result of oxidative stress and chronic inflammation [23]. Oxidative stress is caused by reduced antioxidant levels along with elevated reactive oxygen species (ROS). The proinflammatory cytokine TNF-alpha has been shown to regulate mood and depression [24]. Thus, whether AKE could modulate inflammatory factors and oxidative stress was investigated. To this end, the expression of TNF-alpha, the antioxidant enzyme superoxide dismutase (SOD), and the oxidative damage marker malondialdehyde (MDA) in the brain were measured. These data showed that AKE treatment in male and female rats subjected to overcrowding greatly reduced brain TNF-alpha content (p<0.001). AKE treatment led to increased SOD content, indicative of improved antioxidant status in the brains of male and female rats (p<0.001 for both, Figure 3C). The marker of oxidative stress, total MDA content, was significantly lower both in the brains of male and female rats (p<0.001 for both, Figure 3D). These results revealed that AKE may protect the brain from damage caused by psychosocial stress due to its ability to enhance oxido-inflammatory responses.

## 4. DISCUSSION

Study in 2021 reported that the global prevalence of psychological distress amid the COVID-19 pandemic was estimated at 50.0% [25]. There was a trend towards an increase in psychosocial stress in adolescents [26] and adults [27]. Psychosocial stress has been strongly linked to overall health, yet the intricate processes that connect stress to biological dysfunction are still poorly understood [28]. Various studies seem to suggest the involvement of neuroendocrine and autonomic nervous system abnormalities, inflammation, and oxidative stress [29,30]. Research on animal models of chronic psychosocial stress has demonstrated that behavioral alterations in several tests, such as the open field test and the forced swim test, can mimic the symptoms of clinical anxiety and depression, respectively [31].

The results of the present study showed that twenty-eight days of overcrowding treatment in rats induced a stress-like behavioral and biochemical state, which could be reversed by the AKE. The changes in behavioral and biochemical state of the rats were solely due to overcrowding and AKE treatments, since the environmental conditions for animal housing were controlled at stable condition. The environments were kept stable throughout the experiments with 12 hr light/dark cycle, reduced light intensity (1 - 25 lx), temperature of 22 – 24 °C, humidity of 40-60%, and ad libitum sources of food and water. Many studies have scientifically proven the role of psychosocial stress as a risk factor for depression and depressive symptoms [32]. Indeed, chronic overcrowding treatment in rats in the present study exhibited anxiety- and depressive-related behaviors compared to the pretest. These specific effects were mediated by an alteration in physiological state, including increased corticosterone, decreased testosterone and estrogen, and depleted serotonin (5-HT) levels. Mechanistically, the activation of the HPA axis leads to a surge in corticosterone production [33], resulting in dysregulation of neurotransmitters in the cortical region of the brain, which in turn causes depression-like symptoms [34]. Behavioral changes in stressed rats were also associated with the elevation of MDA, the downregulation of SOD, and the upregulation of TNF-alpha expression in the brain. Co-treatment of overcrowding with AKE attenuated the alterations in behaviors, hormones, neurotransmitters, antioxidants, and cytokine levels elicited by the overcrowding-induced stressful condition.

A proven experimental animal model of psychosocial stress has clearly demonstrated that low levels of brain 5-HT increase stress vulnerability [35]. Hence, most of the current antidepressant medications aim to raise monoamine neurotransmitter concentrations, particularly 5-HT, through selective serotonin reuptake inhibitors. This inhibition is then expected to enhance serotonin signaling [36]. High corticosterone and low 5-HT are the main molecular mechanisms of stress-related depression. A study by Tafet et al. showed that serotonin absorption is increased by heightened cortisol brought on by stress, both at rest and in response to nerve stimulation [37]. In this study, AKE treatment in overcrowding-subjected rats reduced plasma corticosterone levels, which in turn affected serotonergic transmission [31], demonstrating its potent antidepressant activity. The results of the present study are in line with those of previous studies, where antidepressant drugs effectively reduced corticosterone and increased monoamine neurotransmitter levels [38].

In addition to the neuroendocrine systems, there is growing evidence for the involvement of inflammation and oxidative stress in psychosocial stress-related pathology [30,39]. The expression levels of pro-inflammatory cytokines (e.g., IL-1β, TNF-alpha, IL-6, and IL-8) and anti-inflammatory cytokines (e.g., IL-4 and IL-10) in brain tissue were upregulated and downregulated, respectively, by psychosocial stress [40,41]. In the present study, AKE-treated stressed rats showed a lower brain TNF-alpha level, suggesting that AKE prevents brain inflammation induced by overcrowding. The mechanism of the anti-inflammatory property of AKE was revealed by Lee et al. using the RAW 264.7 cell line, which involves the suppression of mitogen-activated protein kinases (MAPK) and nuclear factor-kappaB activation (NF-kB) pathways [42]. The anti-inflammatory activity of AKE is mainly attributed to its flavonoid content [43].

Long-term exposure to psychosocial stress can induce enormous reactive oxygen species (ROS) and oxidative damage, triggering various cellular responses such as neurodegeneration, tissue damage, and cell death that are responsible for the depressive symptoms [12,44]. A study reported that chronic psychosocial stress causes redox perturbations in the brain tissue of rats [45]. In the present study, MDA and SOD were used as oxidative stress markers. MDA is produced as a byproduct of lipid peroxidation, and SOD acts as a radical scavenger by catalyzing the dismutation of the superoxide anion radical into non-reactive molecules [46,47]. Studies showed that the animal model of psychosocial stress exhibits elevated MDA levels and reduced levels of SOD in the brain tissue [48], causing the depletion of 5-HT levels and triggering anxiety- and depressive-related behaviors, as evidenced by the open field and forced swim tests. The AKE treatment in the present study significantly decreased brain MDA levels, accompanied by elevated brain SOD levels, demonstrating the antioxidant properties of AKE. The results of the present study are in line with those of previous studies that showed the beneficial effects of natural antioxidants to attenuate anxiogenic and depressogenic effects [49,50]. A recent meta-analysis study revealed a correlation between the consumption of antioxidant supplements and enhanced states of anxiety and depression, validating their potential as a therapeutic complement to traditional antidepressants [51].

The AKE used in the present study contained 11523.66 mg QE-Eq/g flavonoids, 3100.41 mg GAE-Eq/g polyphenols, 10569.44 mg TAE-Eq/g tannins, an antioxidant capacity of 28294 mg/L GAEAC, and an IC 50% of 80.16 mg/L. A database of species-metabolites called KNApSAcK (http://www.knapsackfamily.com/KNApSAcK/) identified at least 12 types of bioactive compounds in *Angelica keiskei*, including archangelicin, deoxydihydroxanthoangelol H, isobavachalcone, isobavachin, isopimpinellin, osthenol, xanthoangelol, xanthokeismin, xanthotoxin, 4-Hydroxyderricin, and cynaroside. An *in vitro* study by Kim et al. demonstrated that three of the components of AKE (xanthoangelol, 4-hydroxyderricin, and cynaroside) possess antidepressant activity by inhibiting the activity of monoamine oxidases (MAOs), the enzyme responsible for the degradation of amine neurotransmitters, and dopamine β-hydroxylase (DBH). They found that xanthoangelol acts as a nonselective MAO (MAO-A and MAO-B) and DBH inhibitor, 4-hydroxyderricin is a selective MAO-B inhibitor, and cynaroside is a selective DBH inhibitor [16].

In addition to AKE, many medicinal plant sources have been documented to ameliorate the negative impacts of anxiety and depression [15]. The *Caryophyllus aromaticus* extract with a total flavonoid content of 9381.70 mg QE-Eq/g provides antidepressant effects in albino rats [52]. *Sargassum horneri* and *Zataria multiflora* extracts have also been shown to reduce depressive-like behaviors in animal models of stress [53,54]. Lychee peel extract with 90.05 ± 0.26 mg QE-Eq/g total flavonoid content reduces depressive-like behavior in chronic restraint stress-subjected rats [48]. In addition, supplementation of Javanese chili extract (*Piper longum*) with 327.20 mg QE-Eq/g total flavonoid content is also capable of providing antidepressant effects [55]. Rosemary and green tea extracts containing total flavonoid levels of 9075 QE-Eq/g and 605.48 QE-Eq/g, respectively, have been shown to reduce corticosterone levels in stressed animals [56,57]. These reports highlight the potential of natural products with a wide variety of active compounds to treat neuropsychiatric disorders. With the higher content of active compounds (especially flavonoids), AKE is expected to be more effective in preventing and treating anxiety- and depressive-related behaviors and the corresponding biochemical dysregulation. AKE offers a more compelling prospect than synthetic antidepressants and anxiolytics due to the lower toxicity of natural products. It is widely recognized that herbal medicinal products have lower risk for toxicity compared to synthetic drugs [58]. In this study, the side effect of AKE was not observed during experiments at any time point. These findings were also supported by several previous studies that report no adverse side effects for AKE treatment in experimental animal models, and toxicity testing in rats demonstrated that the AKE does not cause mortality or organ damage [59,60]. Despite the promising potential of AKE, further investigation is required to establish the mechanisms of action of AKE as an antidepressant and to perform a thorough toxicological evaluation.

In the present study, control non-stressed animals were not included, which may compromise the conclusion drawn from the results. For the behavioral examinations and hormonal assays, the pretest data were obtained and analyzed, thus the observed effects of overcrowding and AKE treatments could be sufficiently concluded. However, the examination on the brain 5-HT, TNF-alpha, SOD, and MDA levels, where pretest data collection is not possible, cannot be adequately compared because the baseline levels of such substances are not available. This fact limits the conclusion drawn from the brain neurotransmitter, pro-inflammatory cytokine, and oxidative stress markers, as it cannot be discerned as to whether overcrowding causes changes to these parameters and whether AKE treatments reversed them. Although several previous studies have reported that overcrowding induces changes in 5-HT [61,62], TNF-a [63], SOD [64], and MDA [65,66] levels, further study with control non-stressed animals are required to provide more convincing evidence on the role of AKE as anti-depressant by modulating the brain neurotransmitters, pro-inflammatory cytokines, and antioxidant signaling pathways.

## 5. CONCLUSION

In the present study, experimental results show that AKE exhibits antidepressant activities in overcrowding-subjected rats by preventing dysregulation in neuroendocrine, inflammatory, and oxidative stress pathways. The behavioral changes were reversed by AKE, along with lower corticosterone and higher testosterone and estrogen; suggesting that AKE prevents stress-induced infertility and aging-related disorders. In addition, AKE restored the neurotransmitter 5-HT. Furthermore, the SOD was upregulated and MDA and TNF-alpha were downregulated by AKE, demonstrating the potent antioxidant and anti-inflammatory activities of AKE. In conclusion, this study is the first to validate the antidepressant activities of the natural phytomedicine AKE in an animal model, suggesting its potential to be used in clinical settings after clinical trials.

## 8. CONSENT OF ETHICS

All procedures and protocols in this study were approved by the ethical committee of the Faculty of Medicine and Health Sciences, Atma Jaya Catholic University of Indonesia, Jakarta, Indonesia (No. 01/08/KEP-FKIKUAJ/2023).

## 9. DATA AVAILABILITY

All data generated in this study are included in this manuscript.

